# Low-Pass Whole Genome Bisulfite Sequencing of Neonatal Dried Blood Spots Identifies a Role for RUNX1 in Down Syndrome DNA Methylation Profiles

**DOI:** 10.1101/2020.06.18.157693

**Authors:** Benjamin I. Laufer, Hyeyeon Hwang, Julia M. Jianu, Charles E. Mordaunt, Ian F. Korf, Irva Hertz-Picciotto, Janine M. LaSalle

## Abstract

Neonatal dried blood spots (NDBS) are a widely banked sample source that enable retrospective investigation into early-life molecular events. Here, we performed low-pass whole genome bisulfite sequencing (WGBS) of 86 NDBS DNA to examine early-life Down syndrome (DS) DNA methylation profiles. DS represents an example of genetics shaping epigenetics, as multiple array-based studies have demonstrated that trisomy 21 is characterized by genome-wide alterations to DNA methylation. By assaying over 24 million CpG sites, thousands of genome-wide significant (*q* < 0.05) DMRs that distinguished DS from typical development (TD) and idiopathic developmental delay (DD) were identified. Machine learning feature selection refined these DMRs to 22 loci. The DS DMRs mapped to genes involved in neurodevelopment, metabolism, and transcriptional regulation. Based on comparisons to previous DS methylation studies and reference epigenomes, the hypermethylated DS DMRs were significantly (*q* < 0.05) enriched across tissues while the hypomethylated DS DMRs were significantly (*q* < 0.05) enriched for blood-specific chromatin states. A ∼28 kb block of hypermethylation was observed on chromosome 21 in the *RUNX1* locus, which encodes a hematopoietic transcription factor whose binding motif was the most significantly enriched (*q* < 0.05) overall and specifically within the hypomethylated DMRs. Finally, we also identified DMRs that distinguished DS NDBS based on the presence or absence of congenital heart disease (CHD). Together, these results not only demonstrate the utility of low-pass WGBS on NDBS samples for epigenome-wide association studies, but also provide new insights into the early-life mechanisms of epigenomic dysregulation resulting from trisomy 21.

## Introduction

Down Syndrome (DS) is caused by trisomy 21 and is the most common chromosomal aneuploidy present in live births, where it affects approximately 1 in 691.^1^ DS is also the most common genetic cause of intellectual disability and is characterized by distinct facial features and immune system abnormalities. Congenital heart defects (CHD) occur in approximately half of DS patients. Furthermore, DS displays a distinct cancer risk profile, which includes an increased risk for childhood leukemia and a decreased risk for solid tumors.^2,3^ While the genetic basis of DS is well understood, molecular profiling of DS offers insight into the variability in secondary clinical phenotypes, particularly how epigenetic variation can result from a primary genetic change and reflect variable clinical features.

At the molecular level, DS is characterized by differences in gene expression and epigenetic modifications not only on chromosome 21 but also across the entire genome.^4^ Global or region-specific DNA CpG hypermethylation has been observed in a number of DS studies and is most pronounced in the brain and placenta.^5–8^ A meta-analysis of DS DNA methylation array studies uncovered a pan- and multi-tissue CpG methylation signature of DS, where 24 out of the 25 differential genes were hypermethylated in at least 3 of the following tissues: adult brain, fetal brain, placenta, buccal epithelial, and adult blood.^4^ While global hypermethylation (∼1% difference) has been observed in DS adult and fetal brain tissue and sorted cells using arrays, the same study demonstrated that sorted T-lymphocytes (CD3^+^) from adult DS peripheral blood showed the opposite; a trend for global hypomethylation (∼0.3% difference, *p* = 0.186) in DS compared to control.^9^ The sorted DS T-lymphocytes (CD3^+^) showed an approximately equal ratio of hypermethylated and hypomethylated CpGs when compared to controls. Furthermore, the top transcription factor binding site identified within the DS hypomethylated CpGs for the sorted T-lymphocytes (CD3^+^) was the RUNX1 motif.^4,9^ More recently, a separate DS study assayed whole-blood from neonates using array technologies^10^. In this study, differentially methylated regions (DMRs) distinguishing DS from healthy controls showed a slight but significant bias for hypomethylation, although hypermethylation was observed at *RUNX1* (< 1 kb in size). RUNX1 is a developmental transcription factor located on chromosome 21 that is associated with acute myeloid leukemia and classically known to regulate the development of blood cells (hematopoiesis); however, it also regulates neurodevelopment.^11,12^

While reduced representation methods, such as arrays, have provided invaluable insight into the DS methylome, they only assay less than 1 million out of the ∼30 million CpG sites present in the human genome. Previously, in order to assay a larger proportion of the DS methylome, we performed the first whole genome bisulfite sequencing (WGBS) analysis of human DS samples, specifically postmortem brain.^7^ Our low-pass sequencing analysis of post-mortem brains identified 3,152 DMRs, where 75% were hypermethylated and involved in neurodevelopment and metabolism. The hypermethylated DMRs showed a cross-tissue signal, where we replicated many DS pan- and multi-tissue genes and expanded them to larger DMRs, while the hypomethylated DMRs showed a more tissue-specific profile. In this study we present, to our knowledge, the first WGBS of neonatal dried blood spots (NDBS). We assayed more than 24 million CpGs from the NDBS of 86 children of both sexes with either DS, idiopathic developmental delay (DD), or typical development (TD) enrolled in the Childhood Autism Risks from Genetics and Environment (CHARGE) study.^13^ Our analyses demonstrate that the DS newborn blood methylome is distinguishable from that of both TD and DD. We also characterized the profiles of hyper- and hypo-methylated DMRs, which diverge by tissue specificity and suggest a key role for RUNX1 in shaping early-life epigenetic alterations in DS.

## Methods

### Cohort and DNA Extraction from NDBS

Study protocols were approved by the respective Committees for the Protection of Human Subjects at the University of California, Davis and at the State of California. Additional approval was provided by the California Department of Developmental Disabilities and the California Department of Public Health. Informed consent from parents or guardians was obtained prior to collection of any data or specimens. The uploading of the raw or processed sequencing data on individuals to an external bank or repository was prohibited, as the NDBS were property of the Genetic Diseases Screening Program and subject to restrictions in accordance with California Health and Safety Code, Sections 124980(j), 124991 (b), (g), (h) and 103850(a) and (d).

The California Department of Developmental Services’ Regional Center database was used to identify research subjects from the three groups (DS, DD, and TD) within the CHARGE study. The CHARGE participants were matched to the California Newborn Dried Bloodspot Registry, which archives NDBS, also known as Guthrie cards, from heel pricks of newborns obtained 24-72 hours after birth. DNA was extracted from 86 NDBS using protocol GQ, which was established for DNA methylation array profiling of NDBS.^14^ This protocol utilized the GenSolve DNA COMPLETE (GSC-100A, GenTegra) and QIAamp DNA Micro (Qiagen, 56304) kits and was modified to follow the updated manufacturer’s instructions. DNA quality and quantity were assessed via spectrophotometry on a Nanodrop instrument and fluorometry on a Qubit instrument.

### Low-pass WGBS

All DNA samples were sonicated to ∼350bp on a Covaris E220 with a peak power of 175, a duty factor of 10, a cycle/burst of 200, and a time of 47 seconds. A 1.8x SPRI size selection was performed after sonication. Sonication traces were assessed using a Caliper LabChip GX Analyzer. 10 ng of the sonicated and size selected DNA was bisulfite converted using the EZ DNA Methylation-Lightning Kit (Zymo Research, D5031) according to the manufacturer’s instructions. Bisulfite converted DNA was eluted into Low EDTA TE (Swift Biosciences, 90296). Libraries were prepared using the Accel-NGS Methyl-Seq DNA Library Kit (Swift Biosciences, 30096) with the Methyl-Seq Combinatorial Dual Indexing Kit (Swift Biosciences, 38096) according to the manufacturer’s instructions with 12 cycles of indexing PCR. Library traces were assessed using a Caliper LabChip GX Analyzer. The libraries were quantified by fluorometry on a Qubit instrument and pooled in equimolar ratios. The library pool was sequenced across 1 Illumina NovaSeq 6000 S4 flow cell (4 lanes) for 150bp paired end reads to generate ∼100 million read-pairs (∼5X coverage) of the genome per sample.

### Sequencing alignment

Raw sequencing reads were demultiplexed and sample specific FASTQ files were merged across lanes. The FASTQ files were then aligned to the human genome (hg38) using CpG_Me (https://github.com/ben-laufer/CpG_Me) with the default parameters. The pipeline consisted of trimming adapters and methylation bias, screening for contaminating genomes, aligning to the reference genome, removing PCR duplicates, calculating coverage, calculating insert size, extracting CpG methylation, generating a genome-wide cytosine report (CpG count matrix), as well as examining quality control metrics.^15–17^

### Copy number variation (CNV) calling algorithm

A novel read depth based CNV calling algorithm (https://github.com/hyeyeon-hwang/CNV_Me) was utilized to detect large-scale structural variation among the DS, DD, and TD samples. The coverage of 5kb bins in the chromosomes of each sample was calculated by dividing the sum of the number of reads times the read lengths in each bin by the bin size of 5kb. Each bin coverage was then normalized by dividing the sum of the number of reads times read lengths in each bin. To calculate the copy number, the normalized bin coverage value was divided by a normalization factor. To anchor the normal copy number value of autosomal chromosomes to be 2, the normalization factor of autosomes was calculated by multiplying 0.5 by the sum of the normalized bin coverage of all the typical development control samples and dividing by the total number of control samples. The normalization factors of the sex chromosomes were calculated similarly, except the normal copy number of the sex chromosomes was anchored to 1 for male samples. The sex of each sample was verified using k-means clustering on the ratios of the number of reads aligned to the sex chromosomes.

### DMR and block analyses

DMR and block calling as well as some of the downstream enrichment analyses were performed using DMRichR (https://github.com/ben-laufer/DMRichR). To find DMRs with at least a 5% difference, the default parameters were used, aside from directly adjusting for sex, setting perGroup to 0.75, and setting the block permutations to 50. DMRichR utilizes the dmrseq^18^ and bsseq^19^ algorithms to infer methylation levels from CpG count matrices and identify DMRs. These algorithms utilize smoothing and weighting based approaches to infer DMRs from low-pass WGBS, where CpGs with higher coverage are given higher weight. Therefore, the statistical approaches benefit from additional biological replicates more so than deeper sequencing. DMRichR utilizes the dmrseq algorithm to identify DMRs (a few hundred bp to several kb) as well as blocks (> 5 kb), where the coverage filtering settings exceeded the established minimum requirements.^18^ The dmrseq algorithm first assembles candidate background regions, which show a difference between groups, and then performs a statistical analysis to estimate a region statistic. Finally, permutation testing of the pooled null distribution is utilized to identify significant DMRs by calculating empirical *p*-values that are then FDR corrected (*q*-values). Individual smoothed methylation levels for downstream analyses and data visualization were obtained using bsseq.^19^

### Machine learning

Random forest and support vector machine algorithms within the Boruta^20^ and sigFeature^21^ packages, respectively, were used to build binary classification models and generate two lists of the DMRs ranked by variable importance for the feature selection analyses. From the two lists, the common DMRs within the top 1% of each were selected as minimal DMRs. The follow-up machine learning analyses to predict the class of diagnosis from the minimal DMRs identified in the feature selection analyses utilized the random forest algorithm and 5-fold cross validation. The ntree (number of trees) and mtry (number of predictors sampled at each tree split) model parameters were 500 and 2, respectively.

### Enrichment testing

GO enrichment testing was performed using a customized version of GOfuncR, which was based on genomic coordinates and relative to background regions with regions being annotated to a gene if they were between 5 kb upstream to 1 kb downstream of the gene body.^22,23^ The identified significant (*p*_unadjusted_ < 0.05) GO terms were then slimmed using REVIGO and ranked by dispensability.^24^ The hypergeometric optimization of motif enrichment (HOMER) toolset was utilized to test for enriched transcription factor motifs in DMRs relative to background regions through the findMotifsGenome.pl script, where the region size was set to size given and the normalization was set to CpG content.^25^ The genomic association tester (GAT) was utilized to test for sequence specific overlap relative to background regions with GC content correction for the pan- and multi-tissue enrichment testing as well as the genic and CpG annotation enrichment testing.^26^ 10,000 random samplings were used for all GAT analyses. Previous DS datasets were obtained from their respective publications and genic and CpG annotations for hg38 were obtained from annotatr.^27^ The locus overlap analysis (LOLA)^28^ program was also utilized for DMR enrichment testing, relative to background regions, for the reference epigenome histone post-translational modifications (5 marks, 127 epigenomes) and the related chromHMM chromatin states from the core 15-state model.^29,30^

## Results

### Low-pass WGBS of NDBS DNA detects both trisomy 21 and a trend of global hypomethylation

DNA was isolated from 21 DS (13 Male, 8 Female), 33 DD (16 Male, 17 Female), and 32 (16 Male and 16 Female) TD archived NDBS. WGBS sequencing libraries prepared from 10 ng of DNA per sample were indexed and pooled for sequencing across a single NovaSeq 6000 S4 flow cell (4 lanes) to obtain ∼5x coverage (**Supplementary Table 1**). By comparing read depth across each chromosome, a novel copy number variation (CNV) calling algorithm was utilized to confirm trisomy 21 in the DS samples (**Figure 1A**) and to rule out the possibility of other large CNVs on chromosome 21 in the DD and TD samples. Smoothed methylomes from the 24,456,995 CpGs assayed in all groups were first used to detect possible differences in global methylation levels. The DS group showed the lowest global CpG methylation levels (82.1%), followed by DD (82.5%), and TD (82.7%) (**Figure 1B**). When compared to the DD and TD groups, there was a trend (*p*_Diagnosis_ = 0.1, two-way ANOVA) for a slight (∼0.5%) decrease in global CpG methylation levels in DS samples.

**Figure 1:**
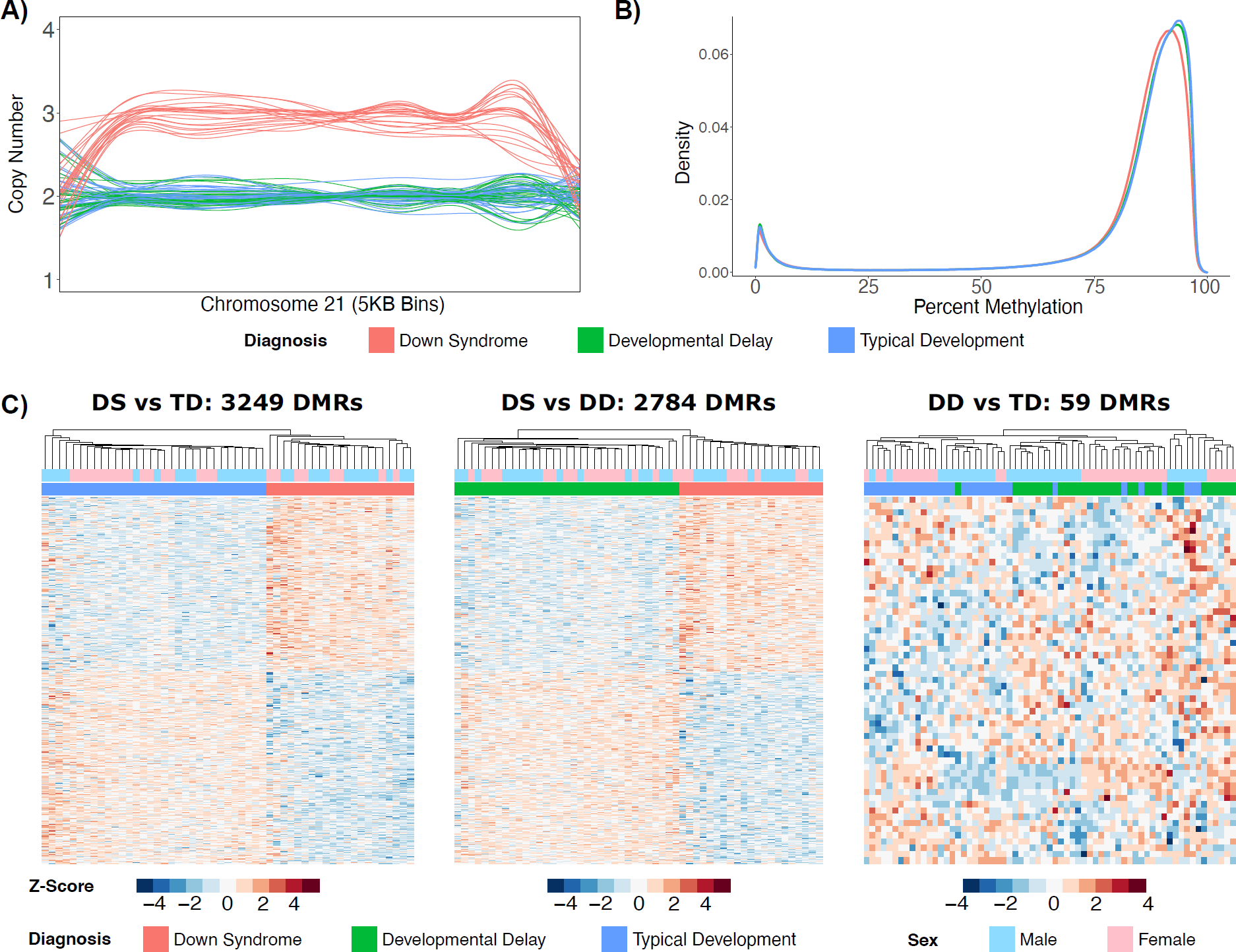
Distinct DS methylome profiles. **A)** Line plot of normalized copy number based on read depth over 5kb bins for chromosome 21 in all 86 samples. **B)** Density plot of average percent smoothed methylation for CpGs covered in the 3 diagnostic groups. **C)** Heatmaps of significant (*q* < 0.05) DMRs from the DS vs TD comparison, significant (*q* < 0.05) DMRs from the DS vs DD comparison, and significant (*p* < 0.05) DMRs from the DD vs TD comparison. All heatmaps display hierarchal clustering of Z-scores, which are the number of standard deviations from the mean of non-adjusted percent smoothed individual methylation values for each DMR.

### Genes associated with DMRs specific to DS reflect differences in neurodevelopment, metabolism, and transcriptional regulation

In order to identify DMRs that distinguished DS from TD and DD, three comparisons with adjustments for sex were performed. The DS vs TD comparison identified 3,249 genome-wide significant (*q* < 0.05) DMRs (47% hypermethylated, 53% hypomethylated) from 22,608 background regions that were assembled from the 24,659,362 CpGs covered in these groups (**Supplementary Table 2A**). The DS vs DD comparison identified 2,784 genome-wide significant (*q* < 0.05) DMRs (48% hypermethylated, 52% hypomethylated) from 20,540 background regions that were assembled from the 24,770,743 CpGs covered in these groups (**Supplementary Table 2B**). In contrast, the DD vs TD comparison only identified 59 nominally significant (*p* < 0.05) DMRs (54% hypermethylated, 46% hypomethylated) from 1861 background regions that were assembled from the 25,122,420 CpGs covered in these groups (**Supplementary Table 2C**). Hierarchal clustering analysis also showed that while DS DMRs clearly distinguished DS from either DD or TD groups, they did not distinguish DD from TD (**Figure 1C**).

Next, we examined the functional relevance of the DS DMRs. DMRs from all comparisons were distributed throughout the genome with no evident preference for a particular chromosome (**Supplementary Figure 1**). DS DMRs were uniquely and significantly (*q* < 0.05) enriched for CpG islands (**Supplementary Figure 2A**) and 5’ untranslated regions (UTRs) (**Supplementary Figure 2B**). DMRs from all comparisons were mapped to genes, where the top significantly (*p* < 0.05, dispensability ≤ 0.25) enriched Gene Ontology (GO) terms were identified (**Figure 2**). Some neurodevelopmental GO terms were enriched in all comparisons (behavior, juxtaparanode region of axon, and GABA-ergic synapse); however, GO terms unique to DS DMRs included some neurodevelopmental terms (autonomic nervous system development and semaphorin receptor complex/activity) as well as terms related to metabolism (choline metabolic process, hemoglobin metabolic process, pyridoxal phosphate binding, and transaminase activity) and transcriptional regulation (nucleosome and DNA-binding transcription factor activity). The GO terms were relatively similar when split by hyper- and hypo-methylation (**Supplementary Figure 3**).

**Figure 2:**
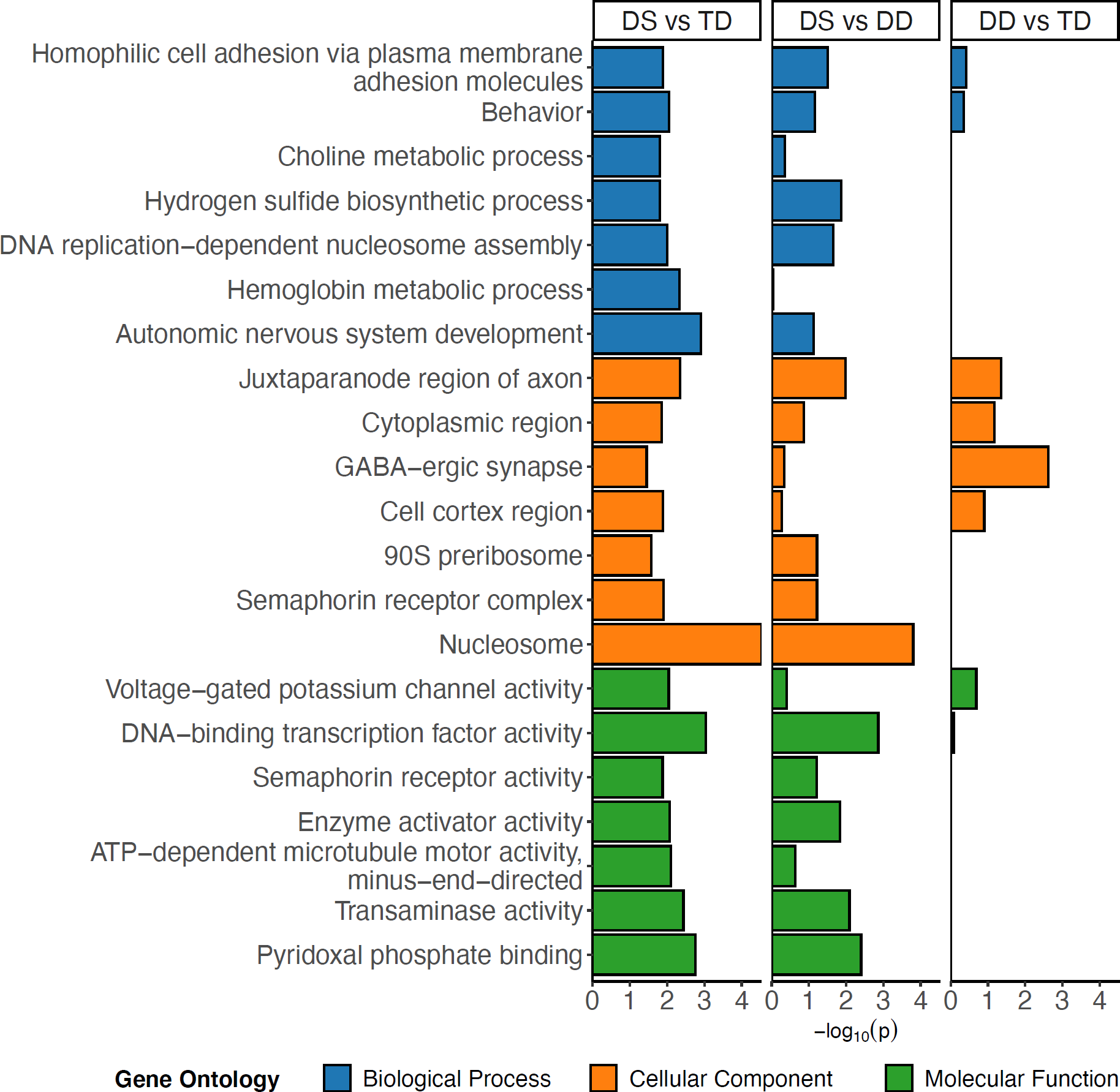
Gene Ontology enrichments. Bar plot of the least dispensable slimmed significant (*p* < 0.05, dispensability ≤ 0.25) GO enrichments for DS vs TD comparison with corresponding values from the DS vs DD and DD vs TD comparison. NAs in the DD vs TD comparison were replaced with 0.

### Consensus DMRs and machine learning predictors distinguish DS from DD and TD NDBS samples

In order to determine if DS DMRs that accurately distinguish DS from idiopathic DD as well as TD samples could be identified, we used two approaches. First, we examined the overlap of DMRs identified in each of the three comparisons (**Figure 3A** and **Supplementary Figure 4A**). Merging the overlaps into single regions that spanned all combined DMRs produced a consensus DMR profile of 4,205 DMRs whose smoothed methylation distinguished DS from DD and TD by the first principle component (**Figure 3B** and **Supplementary Figure 4B**). Second, machine learning feature selection was performed on the consensus DMRs and identified a minimal set of 22 DMRs that distinguished DS from DD and TD (**Figure 3C**). Notably, only one of the DMRs was intergenic and only one was located on chromosome 21 (*RUNX1*). Finally, a machine learning analysis to predict the binary diagnosis class of all 86 samples with the minimal 22 DMRs as predictors performed with an accuracy of 100% and kappa of 1.

**Figure 3:**
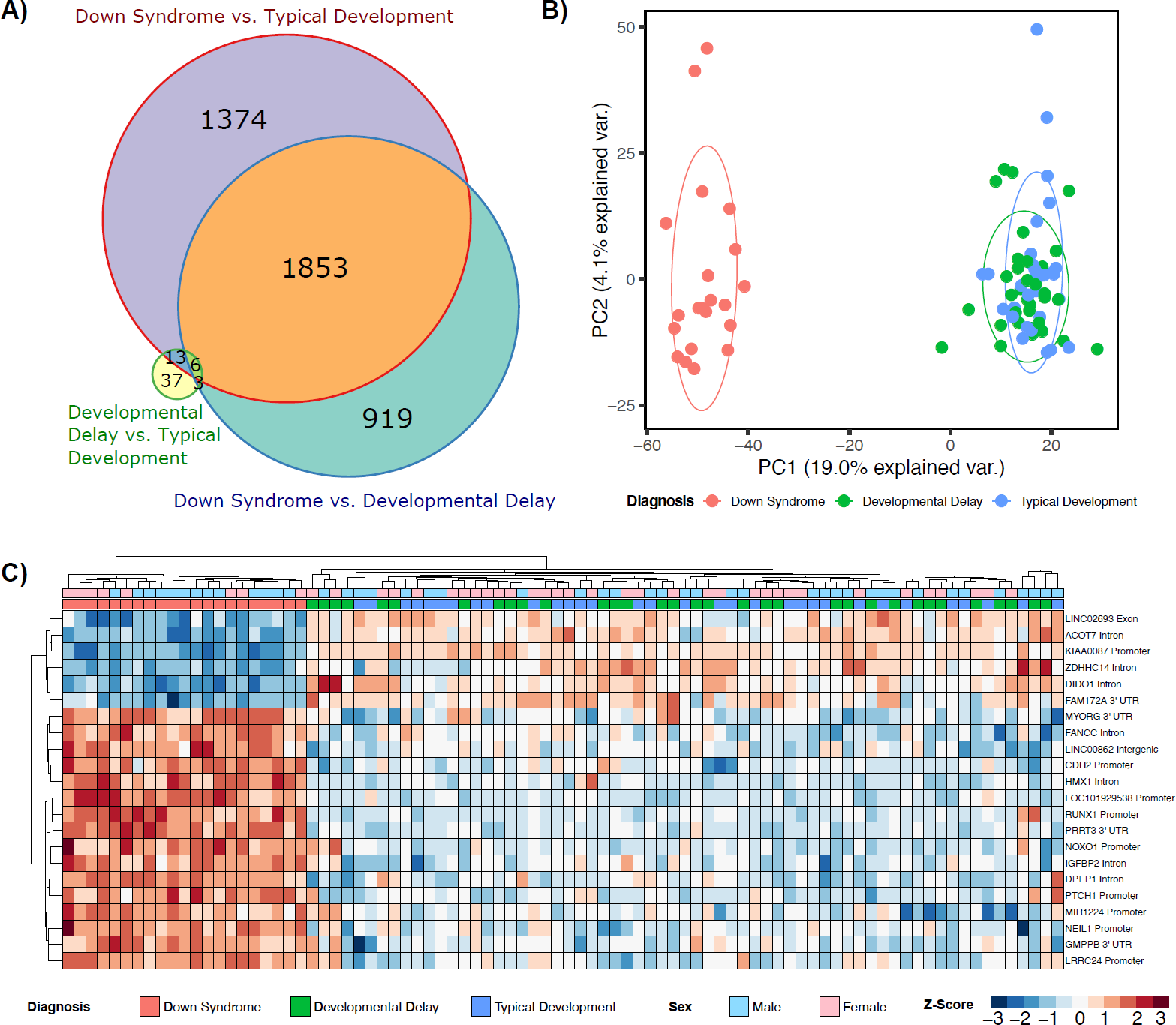
Consensus DMR profiles. **A)** Euler diagram of sequence overlaps for DMRs from all comparisons used to assemble the consensus DMRs. **B)** Principal components analysis of consensus DMRs. Ellipses represent the 68% confidence interval, which is 1 standard deviation from the mean for a normal distribution. **C)** Hierarchal clustering heatmap of the machine learning feature selection analysis of the consensus DMRs.

### Hypermethylated DS DMRs are enriched across multiple tissues while hypomethylated DS DMRs are enriched for blood-specific regions and chromatin states

In order to determine if the DS DMRs we identified in newborn blood are similar to those found in other DS tissues and studies, we performed enrichment analyses separately for hyper- and hypo-methylated DS DMRs for each comparison group (**Figure 4A** and **Supplementary Table 3**). DS hyper- or hypo-methylated CpGs or DMRs were identified from 13 datasets from 8 different studies: 1) Neonatal whole-blood CpGs assayed by Illumina’s Infinium Human Methylation 450K BeadChip array (450K) from Henneman *et al*.^10^, 2) whole-blood CpGs within DMRs assayed on the 450K from Bacalini *et al*.^31^, 3) sorted adult peripheral T-lymphocytes (CD3^+^), adult frontal cortex, sorted adult frontal cortex neurons (NeuN^+^), sorted adult frontal cortex glia (NeuN^-^), adult cerebellar folial cortex, and mid-gestation fetal cerebrum assayed by 450K from Mendioroz *et al.*^9^, 4) buccal epithelium CpGs assayed by 450K from Jones *et al*.^32^, 5) placenta DMRs assayed by Reduced Representation Bisulfite Sequencing (RRBS) from Jin *et al*.^5^ 6) neural induced pluripotent stem cell (iPSC) derivative CpGs assayed by 450K from Laan *et al*.^33^, 7) fetal frontal cortex CpGs assayed by 450K from El Hajj *et al.*^6^, and 8) adult frontal cortex DMRs assayed by WGBS from Laufer *et al*.^7^

**Figure 4:**
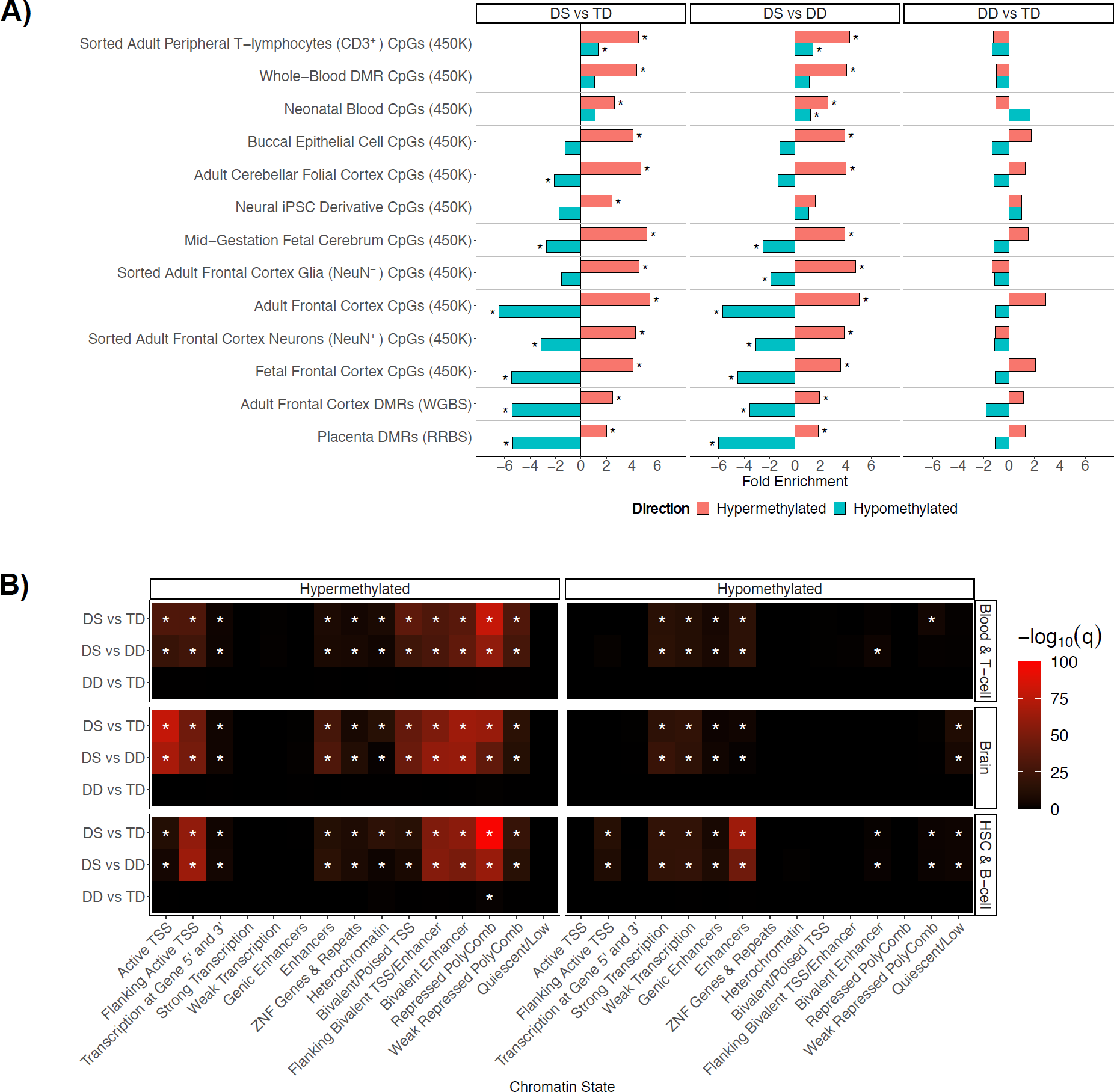
Divergent DNA hyper- and hypo-methylation profiles. **A)** DS cross-tissue enrichments for differential sites from existing DS studies. **B)** Summary heatmap of top q-values for Roadmap epigenomics 127 reference epigenomes chromHMM chromatin state enrichments for all comparisons within the blood and brain tissue reference datasets. * = *q* < 0.05.

As expected, the strongest overall significant (*q* < 0.05) enrichments for the DS NDBS DMRs were within differentially methylated sites from previous DS blood studies, where sorted adult peripheral T-lymphocytes (CD3^+^), whole blood, and neonatal blood were the top ranked. However, in tissues other than blood, the hyper- and hypo-methylated DMRs displayed divergent enrichment profiles with sites from previous DS studies. The DS NDBS hypermethylated DMRs were significantly (*q* < 0.05) enriched for multiple tissues and studies. Additionally, 12 out of the 25 previously known DS pan- and multi-tissue genes were present in our hypermethylated NDBS DMRs, including *RUNX1*, the clustered protocadherins, the *HOXA* cluster, *LRRC24, GLI4, TEX14, RYR1, CYTH2, ZNF837, MZF1, CPT1B*, and *CELSR3*.^4^ In contrast to our hypermethylated DS NDBS DMRs, our hypomethylated DS NDBS DMRs showed significant (*q* < 0.05) de-enrichments in DS differentially methylated sites from non-blood tissues. These results indicate an interesting divergence between hypermethylated regions in DS, which are observed across tissue-type, and hypomethylated regions in DS, which are blood-specific.

To gain further insight into the tissue patterns of DS methylation changes, we performed functional annotation of DMRs using chromatin state segmentations from the chromHMM core 15-state model (based on 5 histone post-translational modifications from 127 reference epigenomes) (**Figure 4B**).^29,30^ DS hypermethylated DMRs showed significant enrichments (*q* < 0.05) across numerous chromatin states, where the strongest enrichments were for repressed polycomb, active transcription start sites (TSS), and bivalent chromatin states. Unlike the hypermethylated DMRs, the hypomethylated DMRs were enriched in a more tissue-specific manner for chromatin states known to vary by tissue type, particularly enhancers in hematopoietic stem cells.

Further investigation of the 5-core histone post-translational modifications from the 127 reference epigenomes also revealed divergence of the hyper- and hypo-methylated DMRs.^30^ Although the top enrichment for the hypermethylated DMRs is from the heterochromatin associated mark H3K9me3 in the thymus (**Supplementary Figure 5A**), which is where T-lymphocytes mature, the hypomethylated DS DMRs in NDBS showed a prominent enrichment for the enhancer associated mark H3K4me1 across blood, and H3K36me3, a mark associated with actively transcribed gene bodies, across multiple cell types (**Supplementary Figure 5B**). Together, these results demonstrate a divergence between the hypermethylated DS DMRs, which contain a cross-tissue signature, and the hypomethylated DS DMRs, which are primarily tissue-specific.

### Early-life RUNX1 dysregulation and hypermethylation is reflected in hypomethylated DS DMRs

In order to identify larger genomic “blocks” of differential methylation in DS, a separate analysis with smoothing parameters optimized to detect regions > 5kb was performed. The DS vs TD analysis identified 3 significant (*q* < 0.05) blocks from 4 background blocks (**Supplementary Table 4A**) and the DS vs DD analysis identified 2 significant (*q* ≤ 0.05) blocks from 3 background blocks (**Supplementary Table 4B**). Two of the previously known pan- and multi-tissue hypermethylated DS DMRs also overlapped within the only significant (*q* ≤ 0.05) blocks in common between the DS vs TD and DS vs DD comparisons, while no blocks were detected in DD vs TD. These two blocks were located in *RUNX1* (**Figure 5A**) and the clustered protocadherins. Overall, *RUNX1* was the highest ranked block and amongst the top two highest ranked DMRs in the DS comparisons.

**Figure 5:**
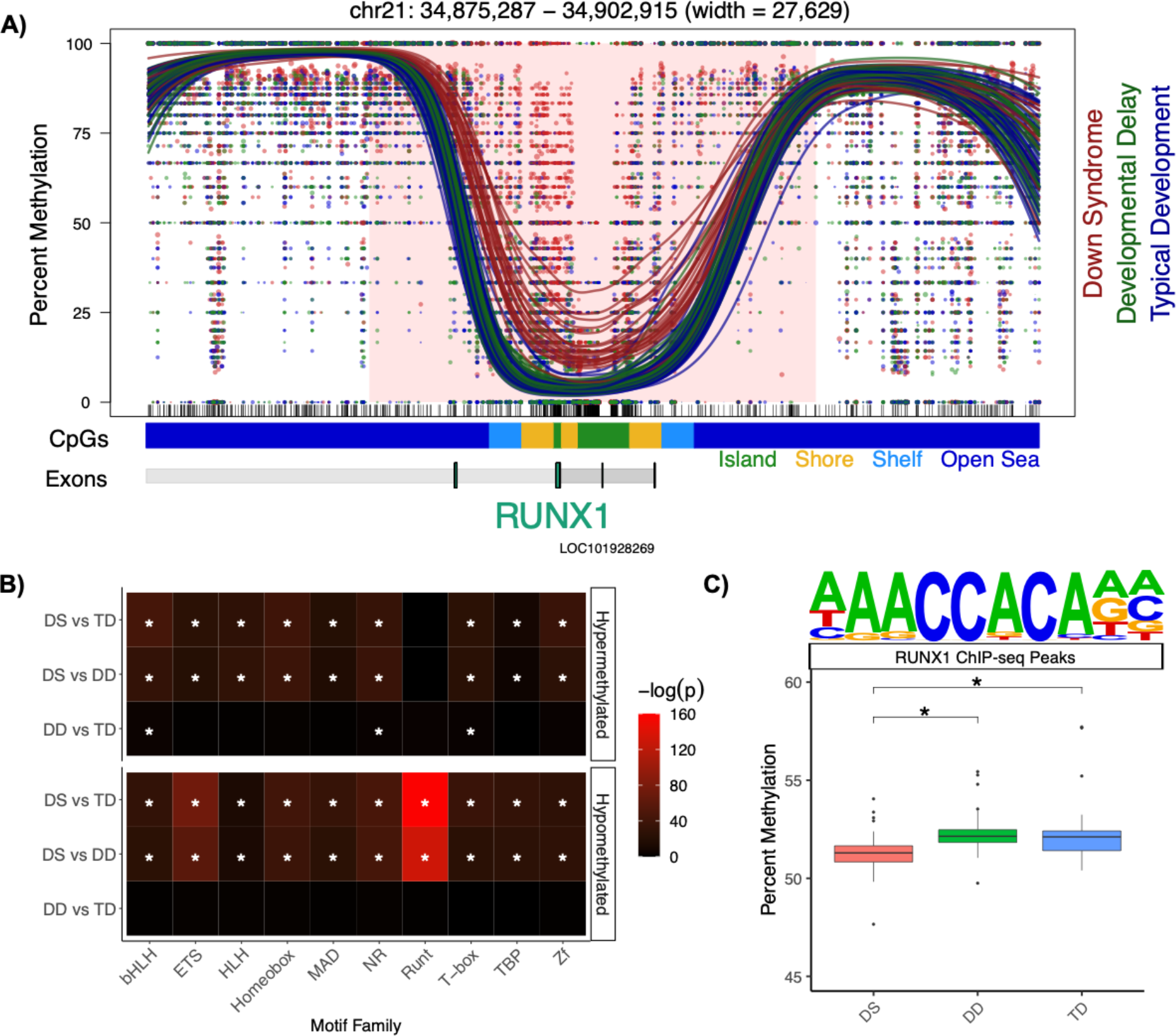
RUNX1 profile. **A)** Significant (*q* < 0.05) hypermethylation within the *RUNX1* block. The lines represent individual smoothed methylation level estimates for DS (red), DD (green), or TD (blue). The dots represent the methylation level estimates of an individual CpG, and the size of each dot is representative of coverage. CpG and genic annotation tracks are shown below each plot, and the *RUNX1* gene is encoded on the negative strand. **B)** Summary heatmap of top *p*-values for top 10 transcription factor motif family enrichments for all comparison groups (* = *q* < 0.05). **C)** Mean percent smoothed DNA methylation levels in RUNX1 binding sites alongside the motif (* = *p*_adjusted_ < 0.05).

An enrichment analysis of transcription factor binding motifs was performed for NDBS DMRs across comparison groups. This analysis identified the Runt motif family as the most significantly (*q* < 0.05) enriched overall, and enrichment for this motif was specific to hypomethylated DS DMRs (**Figure 5B**). The top Runt motif identified belonged to RUNX1 and was from a chromatin immunoprecipitation sequencing (ChIP-seq) experiment that assayed Jurkat cells, which are an immortalized human T-lymphocyte cell line (**Figure 5C**).^34^ A genome-wide analysis, which was not restricted to the DMRs, revealed that diagnosis had a significant (*p* < 0.05, two-way ANOVA) effect on the mean of the smoothed DNA methylation levels within the RUNX1 ChIP-seq peaks. The DS samples had a mean of 51.4% methylation, which was significantly (*p*_adjusted_ < 0.05, post-hoc Tukey HSD tests) lower than the 52.4% methylation of both DD and TD. Taken together, these results indicate that large-scale hypermethylation of RUNX1 (∼ 28 kb) is the strongest signal within the DS DMRs, while its binding sites are specifically associated with the hypomethylated DMR profile as the top transcription factor motif.

### DS patients with CHD display a distinct DMR profile

Amongst the DS cases in our study, there were 11 with CHD (5 Males, 6 Females) and 10 without CHD (8 males, 2 females). In order to identify DMRs that distinguished DS with CHD from DS without CHD, a comparison with an adjustment for sex was performed. There were 1,588 nominally significant (*p* < 0.05) DMRs (35% hypermethylated, 65% hypomethylated) from 50,330 background regions that were assembled from the 22,372,366 CpGs covered in these groups (**Supplementary Table 5**). The DMRs distinguished DS with CHD from DS without CHD (**Supplementary Figure 6A**). Notably, there was an 880 bp hypomethylated DMR that mapped to *RUNX1* in CHD compared to non-CHD DS cases. The only other CHD DMRs that mapped to the known DS pan- and multi-tissue genes were *VPS37B* and the *HOXA* cluster locus, both of which were also hypomethylated, as well as the clustered protocadherin locus, where DMRs distinguishing CHD DS cases were observed in both directions. GO enrichment analyses of the DS CHD distinguishing DMRs revealed significant (*p* < 0.05, dispensability ≤ 0.03) enrichments for terms related to the heart (atrial cardiac muscle tissue development and actin-based cell projection) as well as similar terms to the previous DS vs TD and DS vs DD comparisons, specifically those related to neurodevelopment and metabolism (**Supplementary Figure 6B**). Machine learning feature selection was performed on the DS CHD DMRs and identifed a minimal set of 7 DMRs that distinguished DS with CHD from DS without CHD (**Supplementary Figure 6C**). Additionally, a final machine learning analysis to predict the diagnosis class of all 21 samples with the 7 minimal DMRs as predictors performed with an accuracy of 100% and kappa of 1. The top overall significant (*q* < 0.05) transcription factor motif enrichment for the DS CHD DMRs was for the homeobox transcription factor BAPX1, which was specific to the hypomethylated DMRs (**Supplementary Figure 6D**).^35^ Notably, an ETS:RUNX motif, which refers to regions co-occupied by the two factors, was enriched within the hypermethylated DS CHD DMRs and also within the hypomethylated DS DMRs from the previous comparisons.^36^

## Discussion

Overall, the results of the first WGBS analysis of NDBS DNA samples demonstrate that the DS NDBS methylome is characterized by a genome-wide profile consisting of thousands of DMRs that distinguish it from not only TD but also, for the first time, idiopathic DD. The DS DMRs mapped to genes that are enriched for processes related to neurodevelopment, metabolism, and transcriptional regulation. Furthermore, the hyper- and hypo-methylated DMRs distinguishing DS from DD or TD showed divergent profiles, where the hypermethylated DMRs contained a cross-tissue signature and the hypomethylated DMRs reflected a blood-specific profile related to RUNX1 downstream targets.

One limitation of our study was that neonatal whole-blood is a heterogenous mixture of different cell types, which includes nucleated red blood cells, and alterations in cell composition could influence some of the observed differences. Since our study was, to our knowledge, the first WGBS of NDBS, we utilized multiple existing cell composition estimation methods and reference datasets; however, none accurately estimated the expected cell composition or alterations. Therefore, it was not yet possible to correct for cell type composition due to a lack of a combination of appropriate neonatal whole-blood reference datasets and low-pass WGBS cell composition estimation methods. It is our hope that by demonstrating the feasibility of this assay of a low input sample source for epigenome-wide association studies (EWAS) that future research will generate specific neonatal whole-blood reference datasets and low-pass WGBS cell composition estimation methods. However, we note that our analyses replicate many known DS specific differences that have been observed in analyses of purified cell types (T-lymphocytes)^9^ and bulk tissues (whole-blood)^31^ with cell type composition correction, which both showed the strongest enrichments within our DS discriminating DMRs, a result that would not be expected if cell type shifts had a large effect on our analyses.

Notably, the GO analyses are consistent with the literature and replicate known differences in prior DS metabolome and methylome studies. Previous analyses of plasma metabolites at childhood from DS patients in the CHARGE study revealed distinct alterations to methylation metabolism, specifically choline, which are consistent with our enrichment for DS differentially methylated genes involved in choline metabolism and pyridoxal phosphate (the active form of vitamin B6) binding.^37^ Additionally, our findings replicate those of DS differentially methylated genes from whole-blood, which were involved in processes related to hematopoiesis, neurodevelopment, and chromatin.^31^ Finally, the GO terms identified from DMRs discriminating DS cases with CHD reflect not only heart development, but also neurodevelopmental functions, consistent with the observation that DS infants with CHD are known to show an increased severity of neurodevelopmental disabilities.^38,39^

*RUNX1* is a strong candidate for being a primary dysregulated driver of the epigenetic changes in DS blood. The hypermethylation of *RUNX1* is reproducible across multiple blood studies.^9,10,31^ In our study, hypermethylation of *RUNX1* was the strongest DS signal and it was one of the 22 machine learning predictors of DS in NDBS, where it distinguished DS from not only TD but also idiopathic DD. *RUNX1* was also the top overall transcription factor motif, specifically enriched within the hypomethylated DS DMRs but not the hypermethylated DMRs. The enrichment for *RUNX1* binding motifs in hypomethylated sites has been previously reported in DS T-lymphocytes.^4,9^ Mechanistically, RUNX1 binding results in demethylation of the bound region during hematopoietic development through the recruitment of DNA demethylation enzymes (TET2, TET3, TDG, and GADD45).^40^ Since *RUNX1* is located on chromosome 21, it is overexpressed in DS.^4^ Taken together, these observations suggest that the tissue-specific hypomethylated DMRs we observed in DS NDBS are downstream of a primary alteration to *RUNX1*, where they may have been demethylated by excess RUNX1 binding. This mechanism may also contribute to the observed trend of genome-wide hypomethylation, which has also been previously reported in DS T-lymphocytes.^9^ Notably, the block of hypermethylation overlaps with an enhancer in the first intron of *RUNX1*, which regulates *RUNX1* expression in hematopoietic stem cells and is also part of a larger super-enhancer involved in hematopeisis.^41–43^ Ultimately, the observed alterations in *RUNX1* could influence blood cell function and/or composition. Finally, *RUNX1* methylation could also reflect the severity of secondary clinical features, as a smaller region within *RUNX1* was hypomethylated in DS cases with CHD compared to those without.

Taken together, our findings represent one of the largest sample sizes for a DS methylome study, define a DMR profile that distinguishes DS not only from TD but also idiopathic DD, and provide a novel and general framework for the design and analysis of low-pass WGBS to detect genome-wide methylation changes in NDBS that can be applied to other genetic disorders and environmental exposures.

## Acknowledgements

This work was supported by a National Institutes of Health (NIH) grant [3R01ES015359-10S1] to IHP and JML, a Canadian Institutes of Health Research (CIHR) postdoctoral fellowship [MFE-146824] to BIL, a CIHR Banting postdoctoral fellowship [BPF-162684] to BIL, and the UC Davis Intellectual and Developmental Disabilities Research Center (IDDRC) [NIH U54 HD079125]. The sequencing was carried out by the DNA Technologies and Expression Analysis Cores at the UC Davis Genome Center and was supported by a NIH Shared Instrumentation Grant [1S10OD010786-01]. The authors would like to thank Yunin Ludena Rodriguez for assistance with selecting the samples, Emily Kumimoto for preparing the sequencing libraries, and Matthew Settles for advice about some of the bioinformatic approaches.

## Authorship Contributions

JML, IHP, and BIL designed the study. IHP and JML acquired funding. JML and IFK supervised the project. JMJ and BIL performed the DNA extractions. BIL, HH, and CEM performed the bioinformatic analyses. BIL and JML interpreted the results and wrote the manuscript with intellectual contributions and edits from all other authors. All authors reviewed and approved the final manuscript.

---

**Supplementary Figure 1:**
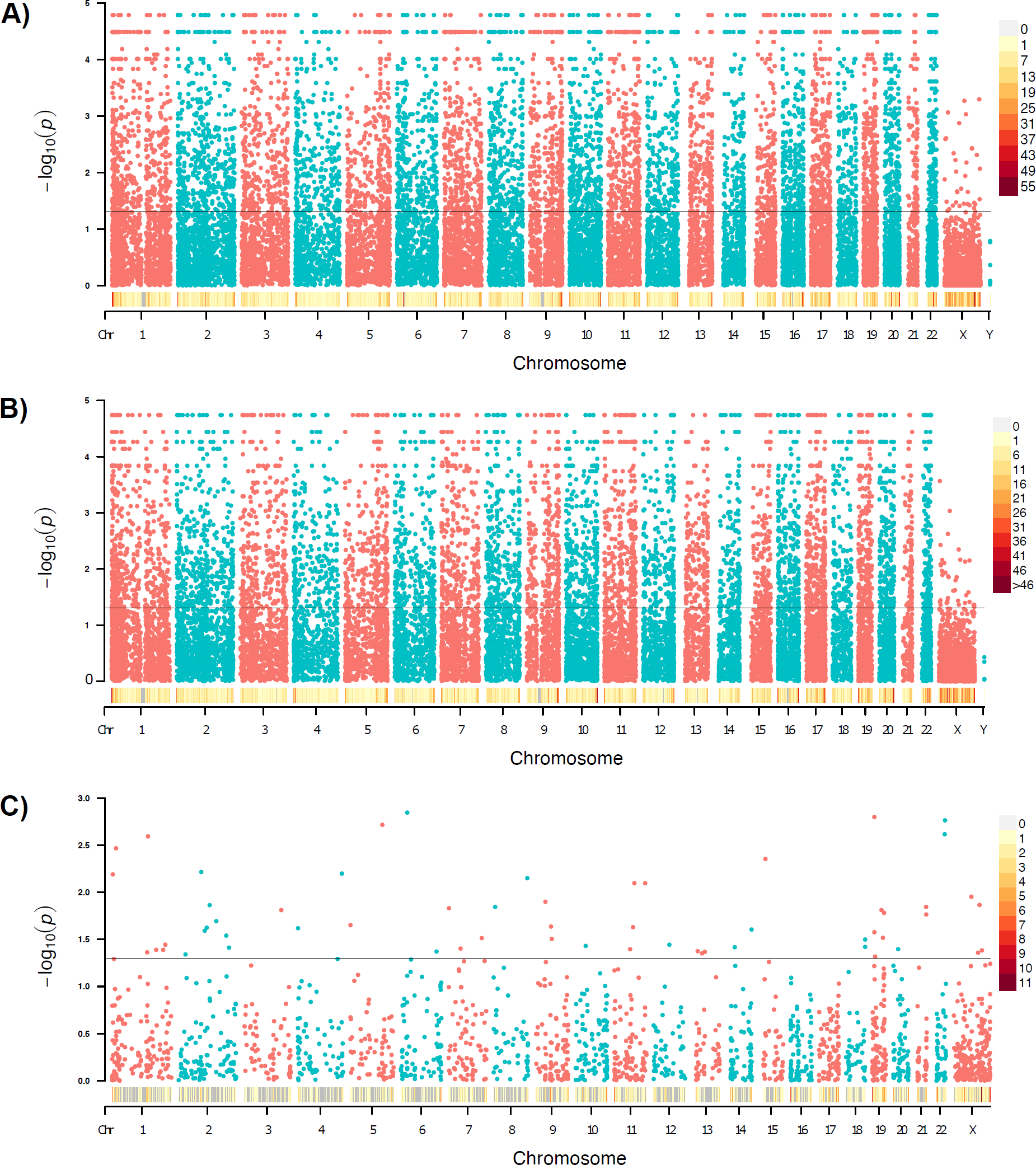
Manhattan plots of tested background regions for the **A)** DS vs TD, **B)** DS vs DD, and **C)** DD vs TD comparisons. The density heatmap indicates the number of DMRs in 1 Mb bins and the line in the Manhattan plot indicates non-adjusted *p* = 0.05.

**Supplementary Figure 2:**
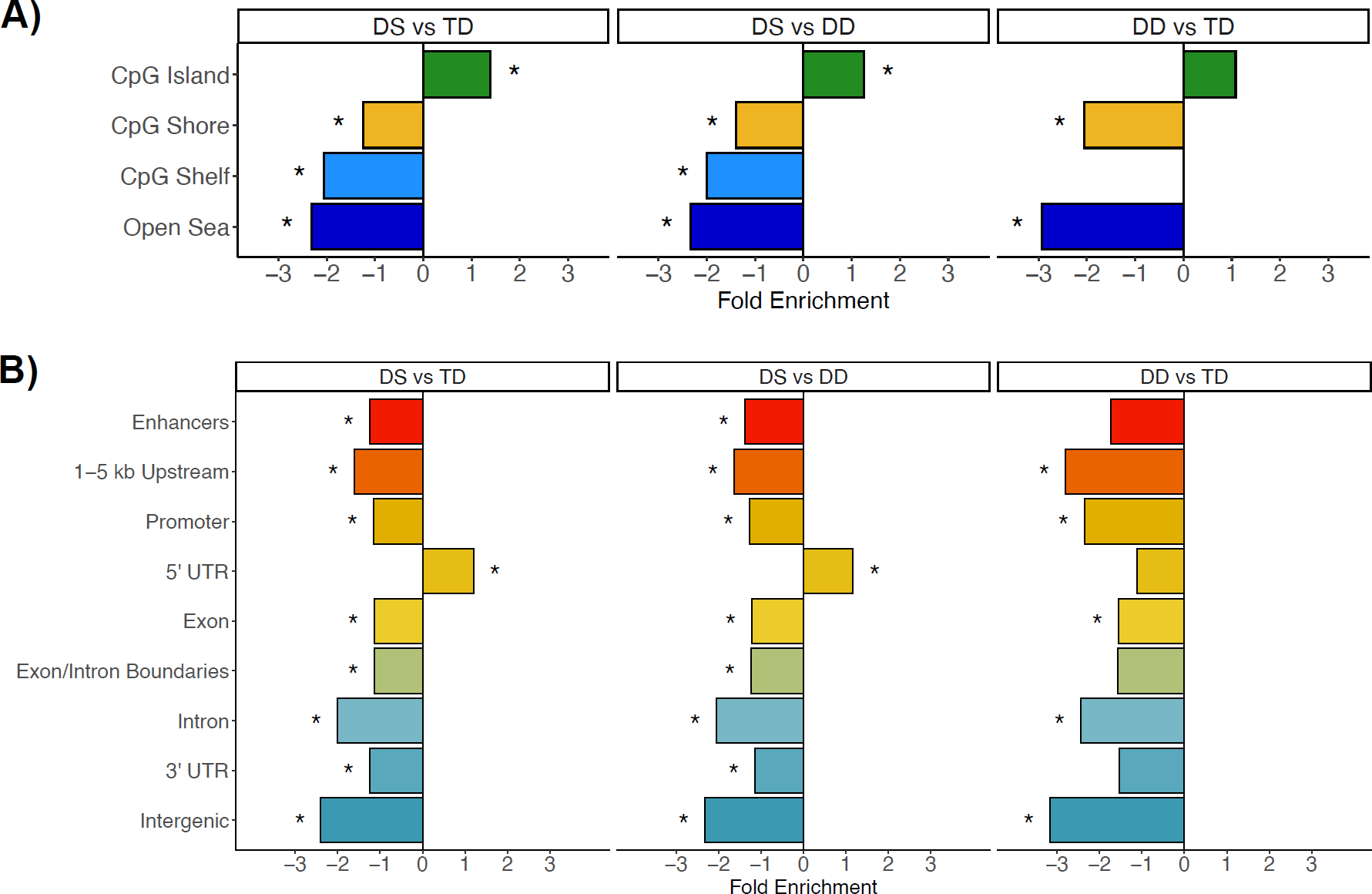
Annotation enrichments for individual DMR comparisons. **A)** Gene region and **B)** CpG annotation enrichments for the DS vs TD, DS vs DD, and DD vs TD comparisons. The enrichments are relative to background regions and corrected for GC content. * = *q* <0.05.

**Supplementary Figure 3:**
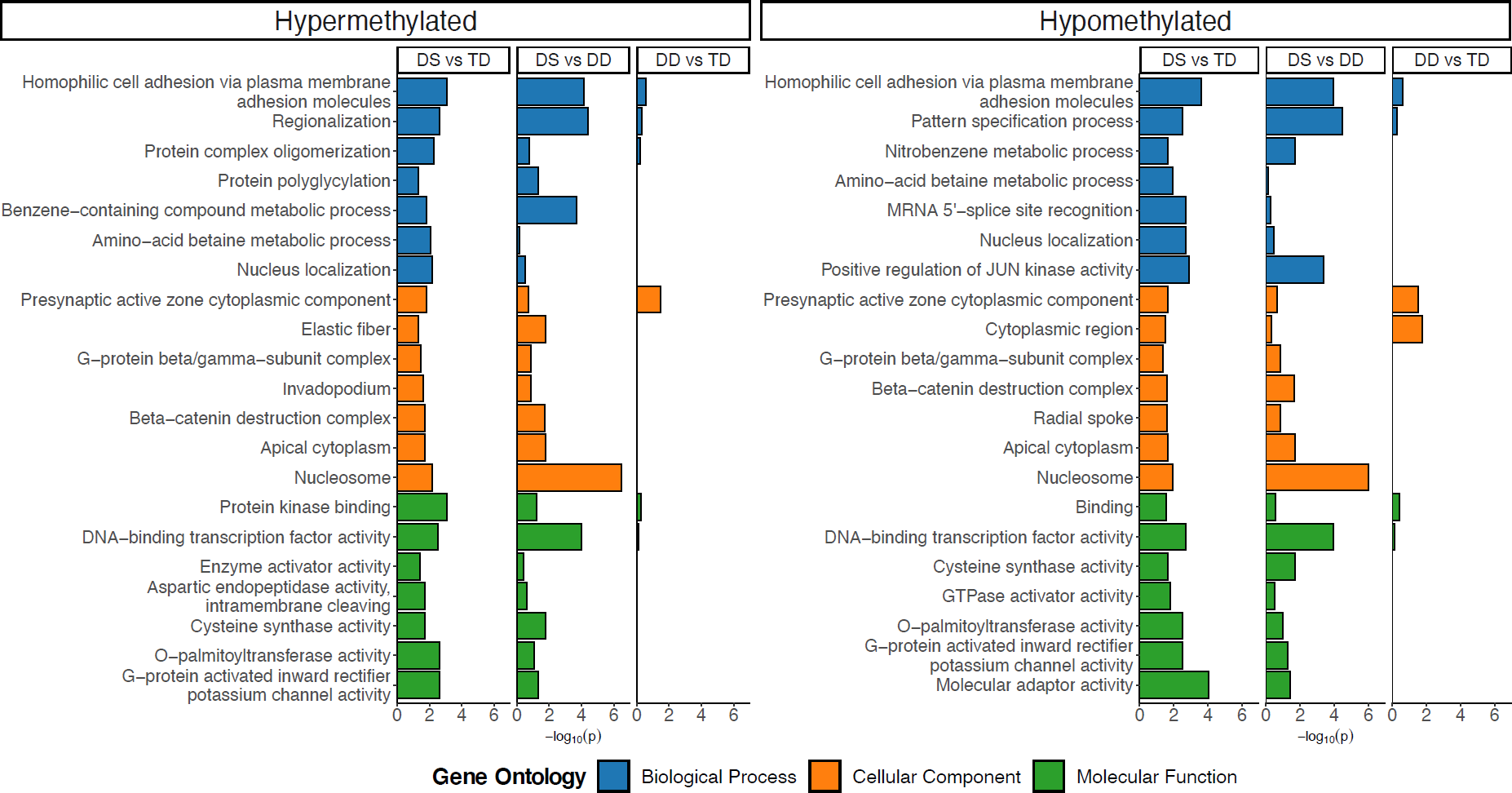
GO enrichments for the hyper- and hypo-methylated DMRs. Bar plot of the least dispensable slimmed significant (*p* < 0.05) GO enrichments for DS vs TD comparison with corresponding values from the DS vs DD and DD vs TD comparison. NAs in the DD vs TD comparison were replaced with 0.

**Supplementary Figure 4:**
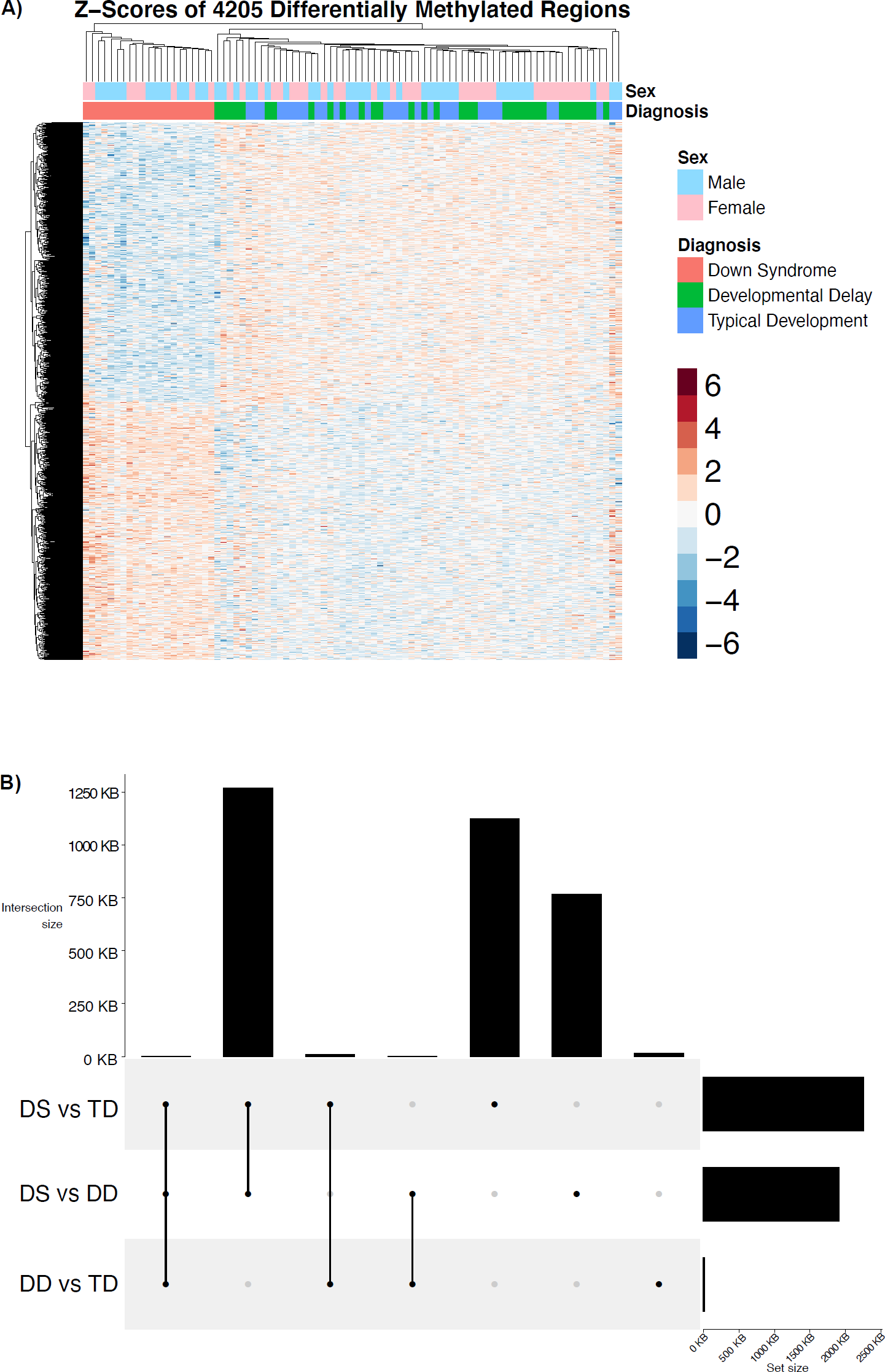
Additional consensus DMR profiles. A) UpSet plot of sequence overlaps for DMRs from all comparisons used to assemble the consensus DMRs. B) Hierarchal clustering heatmap of the consensus DMRs.

**Supplementary Figure 5:**
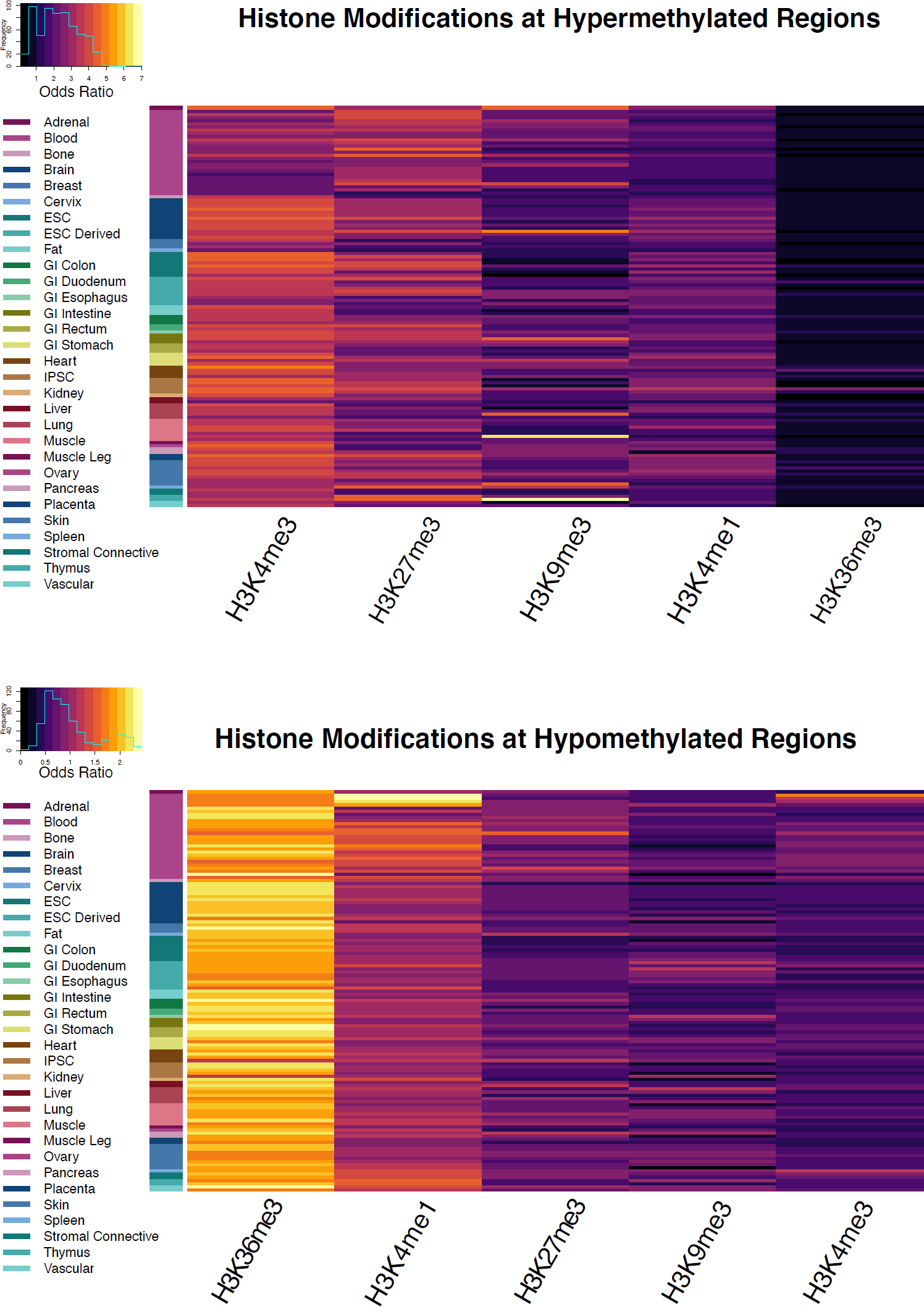
Reference epigenome enrichments for DMRs. **A)** Heatmap of odds ratios for roadmap epigenomics 127 reference epigenomes core histone modifications enrichments for DS vs TD. **B)** Heatmap of p-values for Roadmap epigenomics 127 reference epigenomes chromHMM chromatin state enrichments for all comparisons within the blood and brain tissue reference datasets. All enrichments are relative to background regions. * = *q* < 0.05.

**Supplementary Figure 6:**
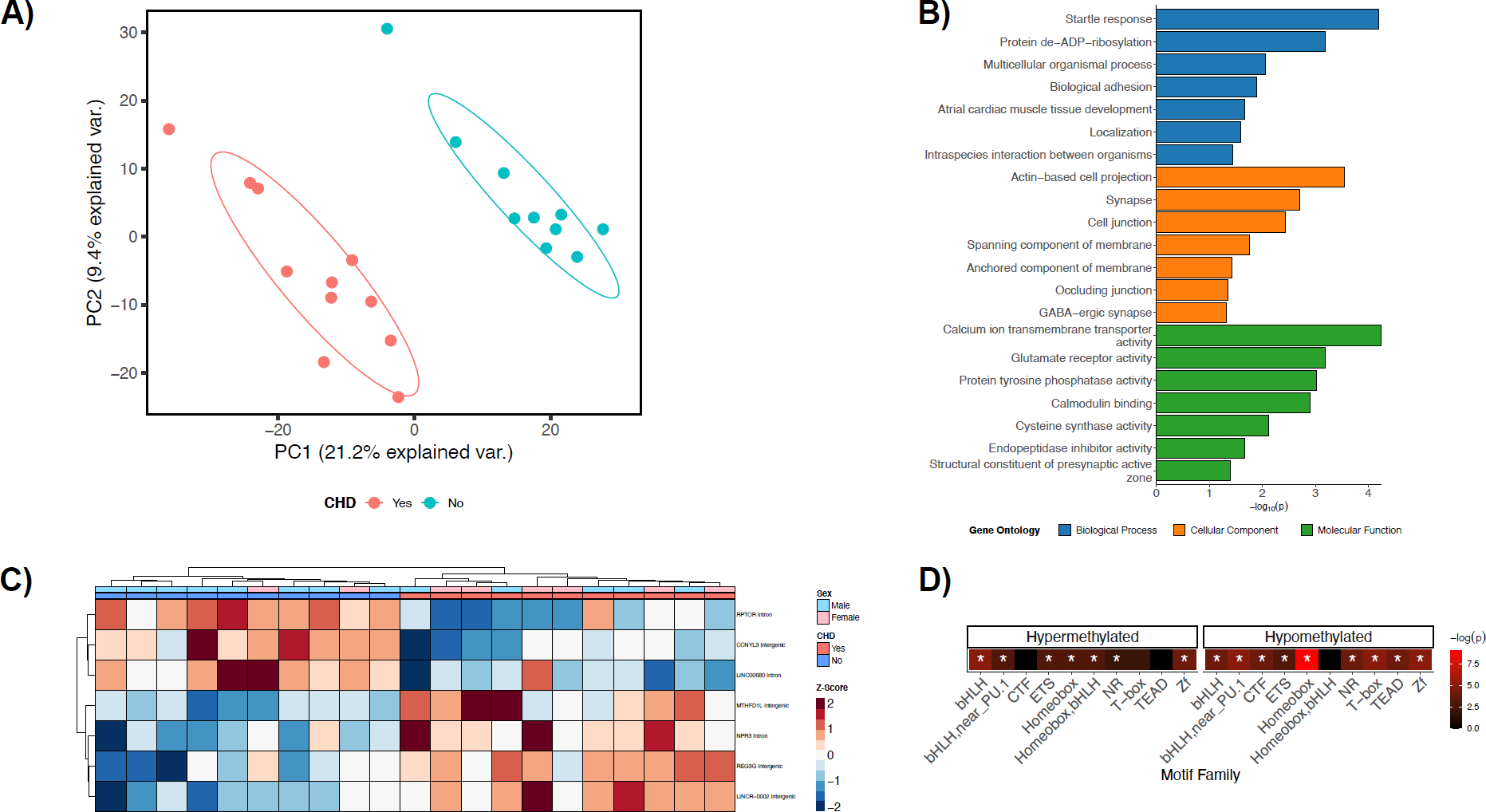
CHD DMR profile. **A)** Principal components analysis of DMRs. Ellipses represent the 68% confidence interval. **B)** Bar plot of the least dispensable slimmed significant (*p* < 0.05, dispensability ≤ 0.03) GO enrichments. **C)** Hierarchal clustering heatmap of the machine learning feature selection analysis. **D)** Summary heatmap of top *p*-values for top 10 transcription factor motif family enrichments (* = q < 0.05).

**Supplementary Table 1**. Subject information and sequencing metrics.

**Supplementary Table 2**. Testable background regions and significant DMRs for **A)** DS vs TD, **B)** DS vs DD, and **C)** DD vs TD.

**Supplementary Table 3:** Previously published DS data sets utilized for overlap enrichment analysis. Reproduced and modified with permission.^7^

**Supplementary Table 4.** Testable background blocks and significant blocks for **A)** DS vs TD and **B)** DS vs DD. There were no testable background blocks DD vs TD.

**Supplementary Table 5.** Testable background regions and significant DMRs for the DS with CHD vs DS with no CHD analysis.

